# Ontogeny-independent expression of LPCAT2 in granuloma macrophages during experimental visceral leishmaniasis

**DOI:** 10.1101/2025.05.23.655708

**Authors:** Shoumit Dey, Jian-Hua Cao, Benjamin Balluff, Nidhi Sharma Dey, Lesley Gilbert, Sally James, Adam A. Dowle, Grant Calder, Peter O’Toole, Ron M. A. Heeren, Paul M. Kaye

## Abstract

Granulomas are organized inflammatory lesions that form in response to persistent stimuli such as infections. Murine infection with *Leishmania donovani* results in the formation of granulomas around infected Kupffer cells in the liver and serves as a well-defined model of immune granuloma formation. The formation and resolution of granulomatous inflammation requires dynamic shifts in immune cell activation states, imposing significant metabolic demands. As mediators of energy homeostasis and cell signaling, lipids and lipid metabolism play a key role in regulating immune cell function during inflammation and the response to infection. However, the extent to which alterations in lipids are spatially linked to altered immune cell transcription has yet to be resolved. In this study, we performed a multimodal imaging analysis combining MALDI mass spectrometry, spatial and single cell transcriptomics, proteomics of flow-sorted macrophages and histopathology of *L. donovani* induced hepatic granulomas. Using this spatially-integrated approach, we identified LPCAT2-mediated membrane re-modelling of myeloid cells as a novel feature of these granulomas. Our study provides new insights into local immunometabolic changes associated with granuloma formation and macrophage activation.

## 1. Introduction

Granuloma formation is a hallmark of many infectious and inflammatory diseases^1^. Well-organized granulomas characterize chronic infections like tuberculosis, where they contain bacteria but also provide a niche for persistence^2,3^. In sarcoidosis, aberrant granuloma responses cause organ damage^4^. Leprosy presents across a spectrum of granulomatous inflammation, with well-formed granulomas correlating with the self-limited tuberculoid form versus poorly organized inflammation in the disseminated lepromatous form^5^. In human and canine visceral leishmaniasis, granulomas are typically absent or poorly formed in clinical cases but demonstrable in asymptomatic individuals^6^, suggesting a host protective role^7^.

In mice, infection with *Leishmania donovani* results in the development of organized hepatic granulomas around infected Kupffer cells. The granulomatous response helps control parasite replication and resolve infection^8^. Surrounding a core of parasitized Kupffer cells other cell types accumulate, including CD4^+^ T cells, CD8^+^ T cells, monocytes, monocyte-derived macrophages (MDMs) and scant neutrophils under the influence of an evolving landscape of cytokine and chemokine expression. Within granulomas, the complex local cytokine environment in part dictates parasite survival or eliminationl^9^, the latter leading over time to granuloma resolution (involution). The formation and resolution of granulomatous inflammation requires dynamic shifts in immune cell populations and functional states, imposing significant metabolic demands. Upon activation, immune cells undergo metabolic reprogramming to support increased energy utilization and biosynthesis^10^.

For example, inflammatory macrophages show increased glycolysis and decreases in TCA cycle and oxidative phosphorylation, and increased lipid metabolism^11^. As mediators of energy homeostasis and cell signaling, lipids and lipid metabolism are intertwined with immune cell activation^12^. In sterile disease, increased cholesterol and fatty acid (FA) biosynthesis, intracellular ceramide, and very-low-density lipoprotein (VLDL) secretion feedback to regulate immune responses^13^. For example, decreased expression of PLA2G7, a lipoprotein-associated phospholipase A2 (Lp-PLA2), was correlated with lower inflammation in calorie-restricted humans and improved metabolism in ageing mice^14^. Lp-PLA2 has also been shown to modulate macrophage M1 polarization in experimental autoimmune encephalitis^15^. These data suggest a microenvironment-specific, direct effect of lipid metabolism on macrophage activation^16,17^. Infectious diseases including TB and leishmaniasis have also been shown to result in altered lipid metabolism^18,19^ though only rarely has this been analyzed in a spatial context ^20,21^.

In this study, we performed multimodal imaging that combines Matrix-assisted laser desorption/ionization mass spectrometry (MALDI) based lipid imaging (MSI), spatial transcriptomics (10x Visium), histopathology of hepatic granulomas (immunostaining) and *ex vivo* proteomics to study the granuloma microenvironment during experimental *L. donovani* infection. Our integrated approach provides new insights into the immunometabolic changes associated with granulomatous inflammation in the hepatic microcosm and evidences the Lands Cycle^22^ enzyme lysophosphatidylcholine acyltransferase 2 (LPCAT2) and its substrates as key components of granulomatous inflammation in liver and macrophage activation *in vivo*.

## 2. Results

### Defining the cellular and molecular landscape of the L. donovani infected liver

Mature granulomas predominate in the liver of C57BL/6 mice infected for 28 days with *L. donovani*, minimizing heterogeneity due to granuloma evolution^23,24^. Using consecutive serial 7μm sections processed for morphology (immunostaining), MSI and 10x Visium, we probed the transcriptomic and lipidomic landscape (Figure **1A**). To overcome the differences in spatial resolution between MSI (20μm) and Visium (55μm), we averaged 4-6 MSI pixels underlying each transcriptomic spot to generate a final integrated histopathological, transcriptomic and lipidomic map of liver niches at 55μm resolution, referred to as a spot (Figure **1A**). Using adjacent liver tissue from the same animals, we produced a combined single-cell atlas of liver cells from infected and naïve mice to allow deconvolution of cell type abundances within spatial spots. Granulomas had typical morphology (Figure **1B**) and were typically composed between 63-71 cells reported as 95% CI of mean (Figure **1C**).

**Figure 1:**
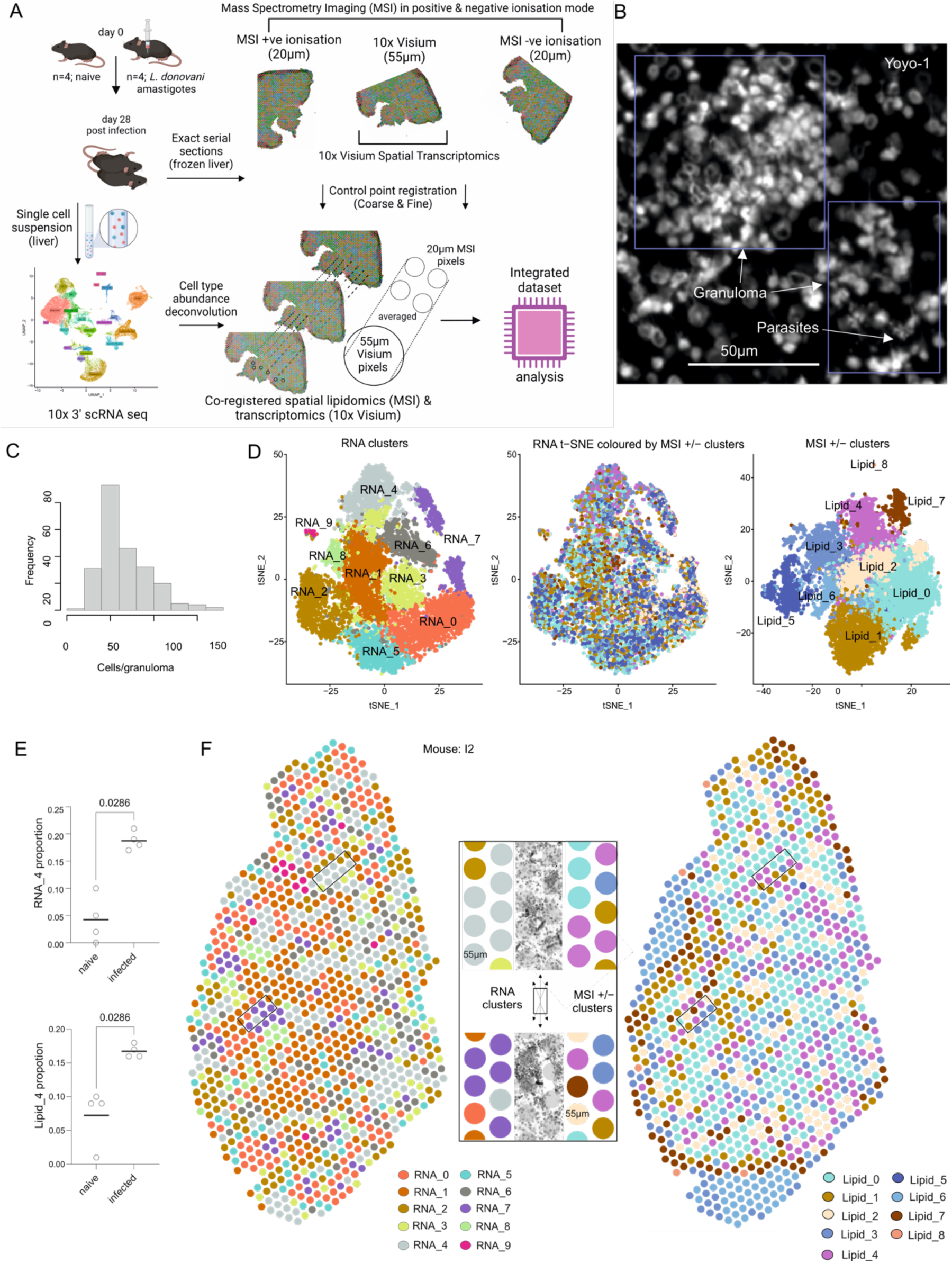
Transcriptomic and lipidomic organization of immune granulomas. **A)** Overall study plan (see Methods for details). **B**) Granuloma morphology using nuclear stain (Yoyo-1, white). Parasites can also be observed in some granulomas. **C**) Distribution of cell numbers per granuloma. **D**) Clusters identified using top principal components separately for RNA and lipids are visualised in t-SNE space (left and right panels, respectively). Spots are visualised in transcriptomic t-SNE space coloured by lipidomic cluster. **E**) Proportional cluster composition of RNA_4 (top) and Lipid_4 (bottom) for naïve and infected mice. P-values are reported using a Mann-Whitney test. **F**) Spatial plot of representative infected tissue showing RNA clusters (left) and Lipid clusters (right). Centre panel shows spots overlying tissue morphology (counterstained by DAPI).

We first performed unbiased clustering of Visium RNA counts and lipid intensities separately (see methods) and visualized these spots in t-SNE space to find underlying structure within the two modalities (Figure **1D**). Upon projecting the lipid-guided clustering information onto the transcriptomics-led t-SNE space, Lipid_4 was found to be enriched within RNA_4 (centre, Figure 1**D**) and both were over-represented in infected mice (Figure **1E** & **S1**). We then demonstrated that our unbiased RNA and lipid clusters showed spatial organization and RNA_4 and Lipid_4 overlaid granulomas defined histologically (**Figure 1F & S1**). In addition, RNA_7 also overlaid some granulomas (Figure **1F**). The top markers of RNA_4 included *Saa3* (Serum amyloid), *Ccl5* and *Lyz2* (Figure **S2A**). Lipid identities (described in **Supplementary Table mz_identifications**) indicated that ether linked PCs such as PC(O-36:3), PC(O-34:2), PC(O-38:5) were observed as the abundant lipid species in Lipid_4 (Figure **S2B**). Overall, we identified spatial cluster organization when independently obtained from both transcriptomic and lipidomic measurements.

### Delineating the Single-cell Transcriptomic Landscape of Granulomas

We performed scRNA-seq on adjacent liver tissue, and manually annotated cell types (Figure **2A****, Supplementary Table scRNA_Celltype_Markers**) to obtain cellular transcriptomes of infected liver non-parenchymal cells. From this, we determined the relative proportions of immune cells across samples (Figure **2B**) using marker genes from imputed clusters (Figure **2C**) and additional canonical markers (Figure **2C**). In naïve mice, cell yields were 7-15×10^6^ and B cells and T cells were represented at similar frequencies. In infected mice, cellularity increased (6-10×10^7^) with *Ifng*^+^CD4^+^ T cells becoming the dominant lymphocyte population (Figure **2B**). We also observed that in infected liver monocyte-derived macrophages (Lyz2Hi_MoMac, positive for *Ccr2*) were nearly twice as frequent than in naïve mice, whereas the proportion of Kupffer cells (ApoeHi_Kupffer, positive for *Adgre1* / F4/80, *Clec4f* and *Marco*) remained similar. Lyz2Hi_MoMac and ApoeHi_Kupffer cells were further distinguished by examining differentially expressed genes, that included some proinflammatory genes such as *Tnf* and *Tlr2* (Figure **2D** **and Figure S3**).

**Figure 2:**
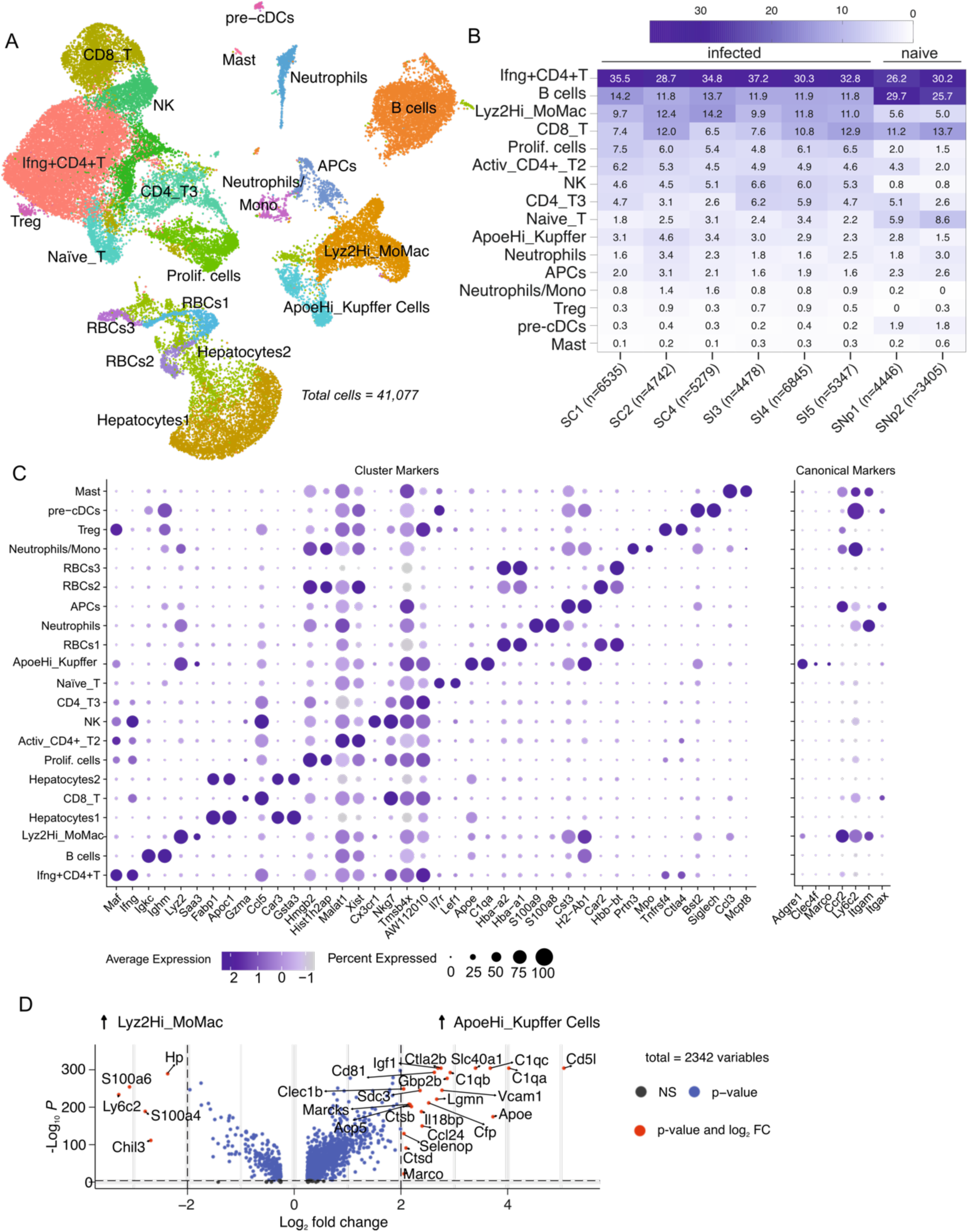
Single-cell transcriptomic landscape of *L. donovani* infected liver. **A**) Imputed cell types visualised in 2-dimensional UMAP space (axes not shown) **B**) Heatmap showing the frequencies of the single-cells across individual infected and naïve samples. Samples names and number of cells sequenced are indicated on x-axis ticks. **C**) Dot plot showing top 2 genes expressed by the imputed cell types. Dot size represents percentage of expressing cells. Colour represents fold change. Canonical markers (right) discriminating cell types. **D**) Volcano plot of differentially expressed genes between Kupffer cells and monocyte-derived macrophages. Bonferroni-corrected Wilcoxon-rank sum test.

### Assessing lipid metabolism pathways in single cells through gene expression patterns

To determine if ApoeHi_Kupffer cells and Lyz2Hi_MoMacs had altered lipid metabolism, we compared the top macrophage upregulated genes (both) with the Reactome (Metabolism of Lipids) gene list for *Mus musculus* (R-MMU-556833). 16 genes were common between ApoeHi_Kupffer cells and Reactome (Figure **3A**), including phospholipases or PLA_2_ that convert phosphatidylcholines (PCs) to lysophosphatidylcholines (LPCs) such as *Pla2g4a* and *Pla2g15.* Additionally, genes downstream to arachidonic acid (AA) metabolism and involved in prostaglandin synthesis (*Ptgs1, Tbxas1*) via the metabolism of polyunsaturated fatty acids (PUFAs) were highly expressed^25^. *Lpcat2* which can incorporate acyl-CoAs to LPCs forming PCs (via the Lands Cycle) was also expressed in ApoeHi_Kupffer cells (Figure **3B**).

**Figure 3:**
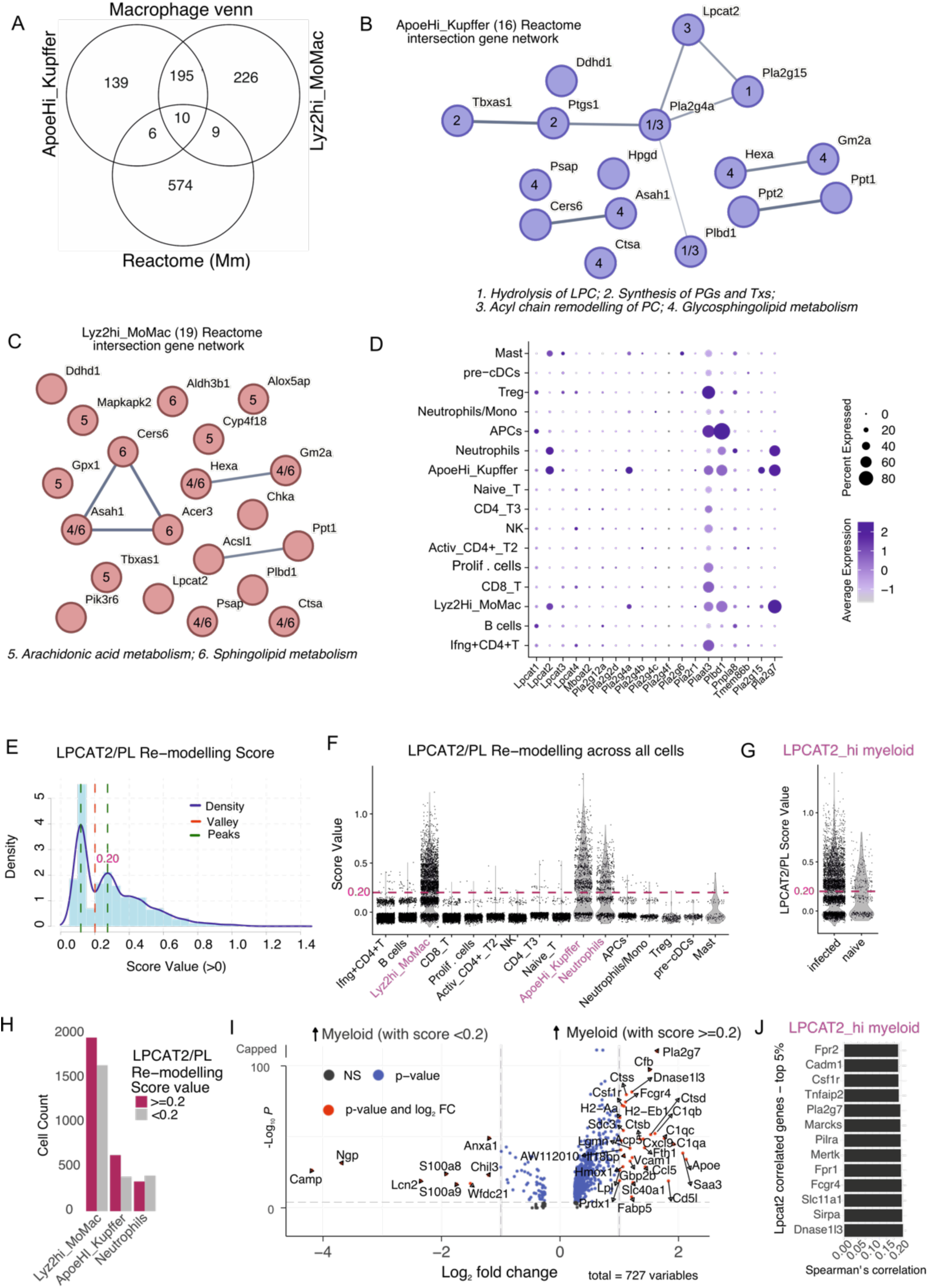
Lands Cycle enzyme LPCAT2 expression in resident and recruited macrophages. **A)** Venn diagram showing the intersection of Reactome Lipid metabolism genes for Mus musculus with top genes expressed by resident Kupffer and monocyte-derived macrophages (MDMs). Genes identified in A are presented as a network of interactions as on STRINGDB and numbered by their role in various reactome pathways as indicated for Kupffer cells (**B**) and MDMs (**C**). **D**) Dot plot showing the expression of Lands Cycle related genes across single-cell populations. **E**) Histogram showing the spread of module score LPCAT2/PL remodelling score (*Lpcat2*, *Pla2g4*, *Pla2g7*) for all cells. Score in E is plotted for each single-cell population (**F**) and for naïve versus infected groups (**G**). **H**) Barplot showing split between populations of Kupffer cells, MDMs and Neutrophils scoring high/low for LPCAT2/PL remodelling. **I**) Volcano plot showing genes upregulated myeloid cells in H that score high Vs. low. **J**) Barplot showing Spearman’s correlation between *Lpcat2* and other genes in high-scoring myeloid cells as in I.

Similarly, the Lyz2Hi_MoMac and Reactome intersection (19 genes; Figure **3A**), also suggested enrichment of sphingolipid and AA metabolism pathways, along with the PC re-modeller *Lpcat2*. As PC hydrolysis can lead to AA release and LPCATs can incorporate acyl-CoAs such arachidonyl-CoA into PCs^26^, we compared acyl-chain remodelling genes across cell types. Both ApoeHi_Kupffer cells and Lyz2Hi_MoMacs upregulated *Lpcat2*, *Pla2g4a*, *Pla2g15* and *Pla2g7*. To confirm whether LPCAT2-mediated PC remodelling occurred on macrophages independent of their origin (*Ccr2*^hi^ or *Adgre1*^hi^) we created a module score for each cell using *Lpcat2*, *Pla2g4a*, *Pla2g15* and *Pla2g7* (Figure **3E**). Highest scores were associated with the two macrophage populations and with neutrophils (Figure **3F**). For these three myeloid cell populations, high scores were associated with infected mice (Figure **3G**) and were observed in ∼50% of each cell type (Figure **3H**). In contrast to more mature macrophages, *Ly6c2*^hi^ neutrophils / monocytes and *Itgax*^+^ APCs had a very low LPCAT2 remodeling scores (Figure **3F**).

Further, there was a change in gene expression between high and low scoring cells, suggesting the former are more activated and/or inflammatory based on the expression of *C1qa*-*c*, MHC-II genes (*H2-Aa* and *H2-Eb1*), *Apoe*, *Cxcl9*, *Saa3* and *Cd5l* (Figure **3I**). Indeed when LPCAT2/PL Remodelling score is correlated against an inflammatory (based on *Tnf*, *Nos2*, *Il6*, *Cd86* and *Tlr2*) or anti-inflammatory (based on *Il10*, *Tgfb1*, *Arg1*, *Il4*) we find remodelling score to correlate higher with the inflammatory score (Figure **S4**). Within myeloid cells with a high LPCAT2/PL remodelling score, *Lpcat2* positively correlated with genes related to phagocytosis, lipid processing, and immune / inflammatory responses, including *Pla2g7*, *Marcks, Sirpa*, *Mertk*, and *Fcgr4* (Figure **3J**).

### Distinguishing granulomatous inflammation

We next used cell2location (see methods) to predict cell abundances within each RNA cluster. RNA_4 was enriched in macrophages, CD4^+^ and CD8^+^ T cells, NK cells and neutrophils. In addition, RNA_0 and RNA_7 were also predicted to contain immune cells with T cell sub-types including Tregs and APCs respectively (Figure **4A**). Hepatocytes were most abundant in RNA_5 (hepatocytes1 and hepatocytes2) and in RNA_0 (hepatocytes2) (Figure **4A**). As immune cells were concentrated in RNA_0, RNA_4 and RNA_7 (Figure **4A**) but immune cells were also found in naïve mice (**Figure 2B**), we combined these immune clusters and performed subclustering, identifying 5 sub-clusters (Sub0-4) which separated infected from naïve tissue (Figure **4B**). Overlaying Sub0 and Sub2 on infected tissue demonstrated a spatial distribution supporting their assignment to granulomas (Figure **4C**). From our lipidomics dataset, Sub0 and Sub2 differentially contained major lipid classes including phosphatidylglycerol (PG), phosphatidylserine (PS), phosphatidylethanolamine (PE), sphingomyelin (SM), phosphatidylcholine (PC), ceramide phosphate (CerP), carnitine (CAR), and inositol-containing lipids such as phosphatidylinositol (PI) inositolphophoceramide (IPC). Ether-linked species were identified in both PC (PC(O-)) and PE (PE(O-)) classes, along with diacylglycerols (DG), diacylglyceryl trimethylalanine (DGTA), triacylglycerol (TG), and cholesteryl ester (CE). Several of these lipids contained arachidonyl acid (20:4)-such as CE(20:4;O2) (Figure S5) and PE(O-18:0_20:4), LPE(20:4), PE(20:0_20:4), LPC(20:4), PS(18:0_20:4) in the top lipid species (Supplementary Table subcluster_lipid_markers) discriminating Sub0 and Sub2 from subclusters found in naïve mice. (Figure **S5**).

**Figure 4:**
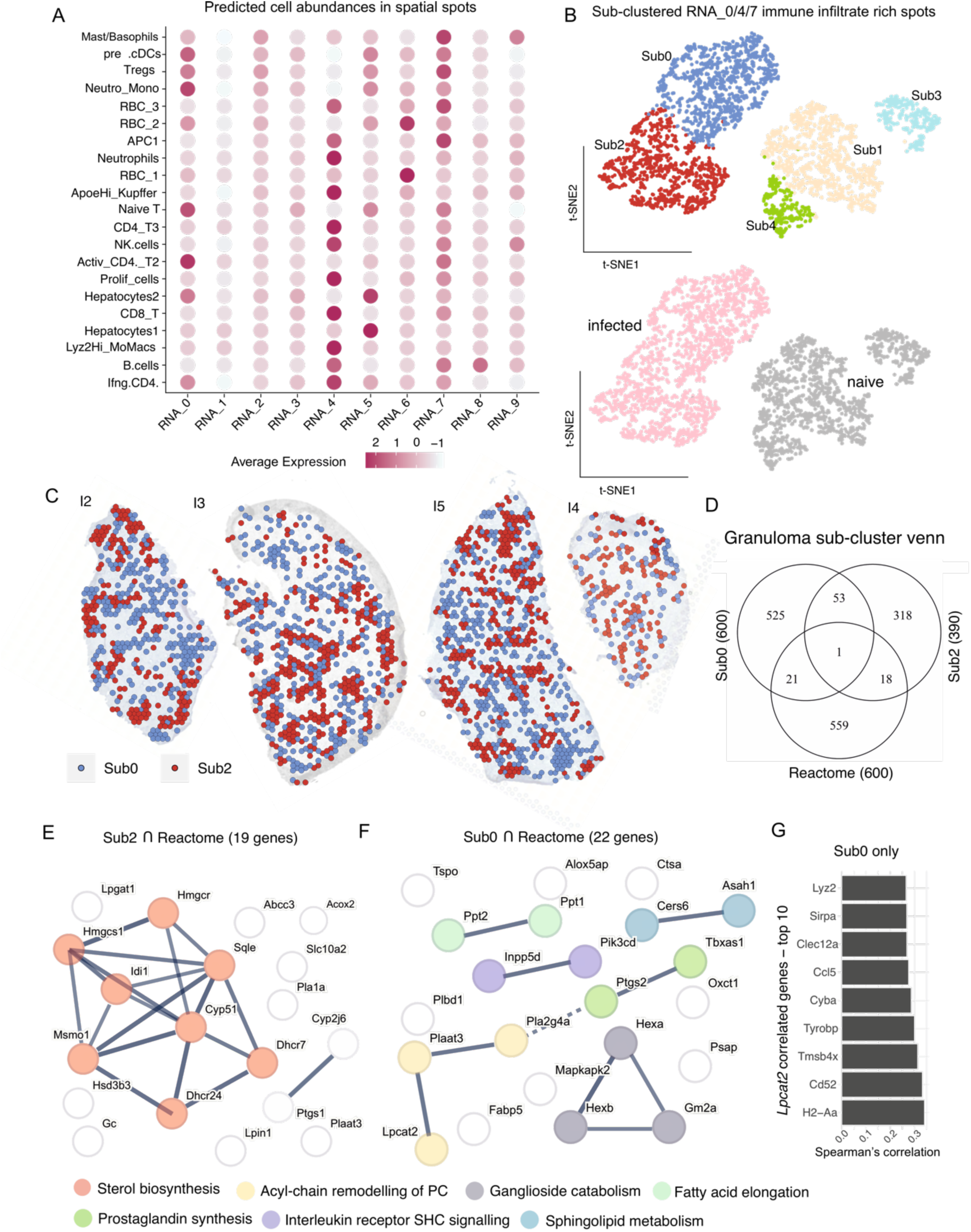
Lipid metabolism related genes in granulomatous clusters. **A**) Dot plot showing predicted cell abundances in RNA_clusters identified in Figure 1. **B**) RNA_0/4/7 re-clustered to identify sub-clusters Sub0-4 (top) associated with infected or naïve mice (bottom). **C**) Spatial representation of Sub0 and Sub2 on infected tissue (n=4 mice). **D**) Venn diagram showing the intersection of Reactome Lipid metabolism genes with top genes expressed by Sub0 and Sub2. **E, F**, STRINGDB network of intersecting genes for Sub2 (E) and Sub0 (F), coloured by MCL clustering and function. Edge thickness represents interaction confidence and dotted lines indicate cluster boundaries. **G.** Spearman’s correlation between *Lpcat2* and genes in Sub0.

Comparing spatial gene expression profiles in Sub0 and Sub2 with Reactome Metabolism of Lipids (Figure **4D**) identified that in sterol biosynthesis was upregulated Sub2 (Figure **4E**), whereas Sub0 suggested enrichment of acyl-chain remodelling of PC. Of note, PG synthesis enzymes such as *Lpcat2*, *Plaat3* and *Pla2g4a* observed in myeloid cells (Figure **3D**) were associated with Sub0 (Figure **4F**). Sub0 also showed enrichment for Sphingolipid metabolism (Figure **4F**). Within Sub0, *Lpcat2* correlated with *H2-Aa*, *CD52*, *Sirpa* amongst others (Figure **4G**).

### Acyl-chain remodelling in granulomas

We next compared the *Lpcat2*, *Pla2g4a, Plaat3, Pla2g15* and *Pla2g7* signature (Figure **5A**) and predicted cellular abundances (Figure **5B**) within Sub0 and Sub2. The identification of *Lpcat3* within Sub0 was consistent with the predicted presence of hepatocytes1^27^ and may reflect some inclusion of hepatocytes in spots at the periphery of granulomas. *Lpcat1* was expressed in Sub0 spots and may reflect lymphocytes composition (Figure **5B**). ApoeHi_Kupffer cells had similar predicted abundance between these subclusters, whereas Lyz2Hi_MoMac were less abundant in Sub2 (Figure **5C**). We further identified differentially expressed genes discriminating these two subclusters (Figure **5D**).

**Figure 5:**
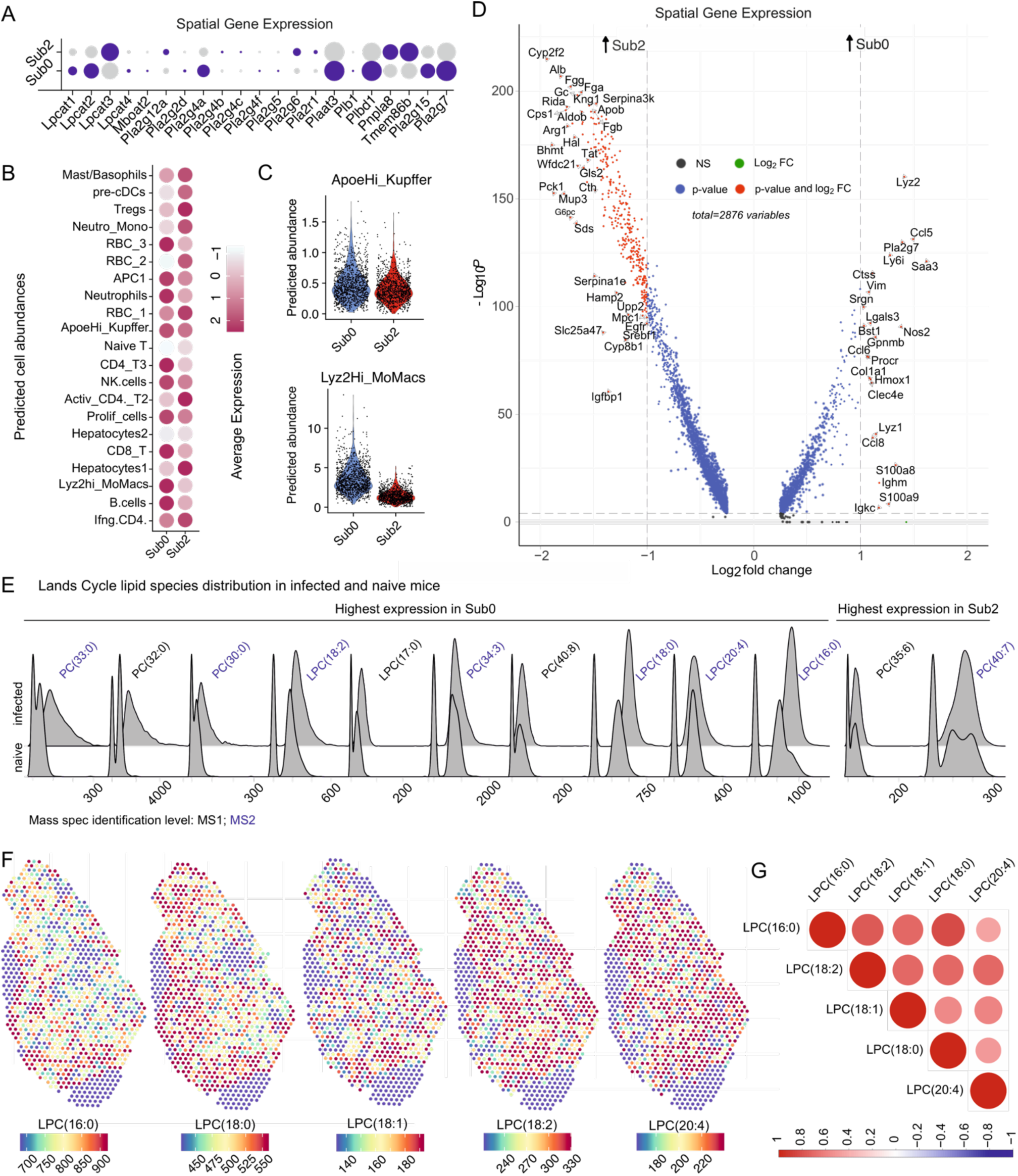
Association of LysoPC species with immune granuloma spots. **A**) Dot plot showing gene expression between Sub0 and Sub2 **B**) Dot plot showing predicted cell abundances in Sub0 and Sub2 **C**) Violin plots showing predicted cell abundances for Kupffer cells and monocyte-derived macrophages **D**) Volcano plot showing genes upregulated in Sub0 versus Sub2 **E**) Ridge plots showing distributions of LysoPCs (LPCs) and PCs with those identified with higher confidence (MS2; data-dependent lipid identification) indicated in blue across infected and naïve mice. **F**) Spatial MSI intensity of LPCs. **G**) Correlation plot (Spearman’s) between LPCs.

To determine whether Sub0 and Sub2 also contained Lands Cycle products and / or substrates for LPCATs, we examined phosphatidylcholines (PCs) identified by MSI. Sub0 showed prominent abundance of PC(30:0), PC(32:0), PC(33:0) PC(34:3), PC(40:8), as well as of all detected lysoPCs LPC(16:0), LPC(17:0), LPC(18:0), LPC(18:2) and LPC(20:4) (Figure **5E**). Notably, LysoPCs showed spatial hotspots (Figure **5F**) and with saturated LPC(18:0) and LPC(16:0) intensities being highly correlated (Figure **5G**). Collectively, these integrated transcriptomic and lipidomic data identify altered expression of Lands cycle enzymes and their associated substrates (and products) within granulomas and implicate ApoeHi_Kupffer cells and Lyz2Hi_MoMacs as the dominant cell types associated with this pathway.

### *Expression of* LPCAT2 *in granulomas*

To validate these findings at the level of protein expression, we stained infected and naïve liver tissue for LPCAT2. LPCAT2 was absent in naïve mice (Figure **6A**) and, in keeping with the mRNA data, LPCAT2 was localized to granulomas in infected mice (Figure **6A-B**). To potentially assign LPCAT2 expression to ApoeHi_Kupffer cells, we co-stained for LPCAT2 with ApoE. Based on dual expression of LPCAT2 and ApoE, approx. 50% of Kupffer cells expressed LPCAT2 (Figure **6D**) ad granulomas contained a similar frequency of other LPCAT2+ cells (likely from our mRNA data to be Lyz2Hi_MoMacs or neutrophils). Only a weak correlation was observed between parasite abundance per granuloma (determined by OPB staining) and the number of LPCAT2+ cells (Figure **6E**), suggesting LPCAT2 expression is driven by local macrophage activation and is largely independent of macrophage infection per se.

**Figure 6:**
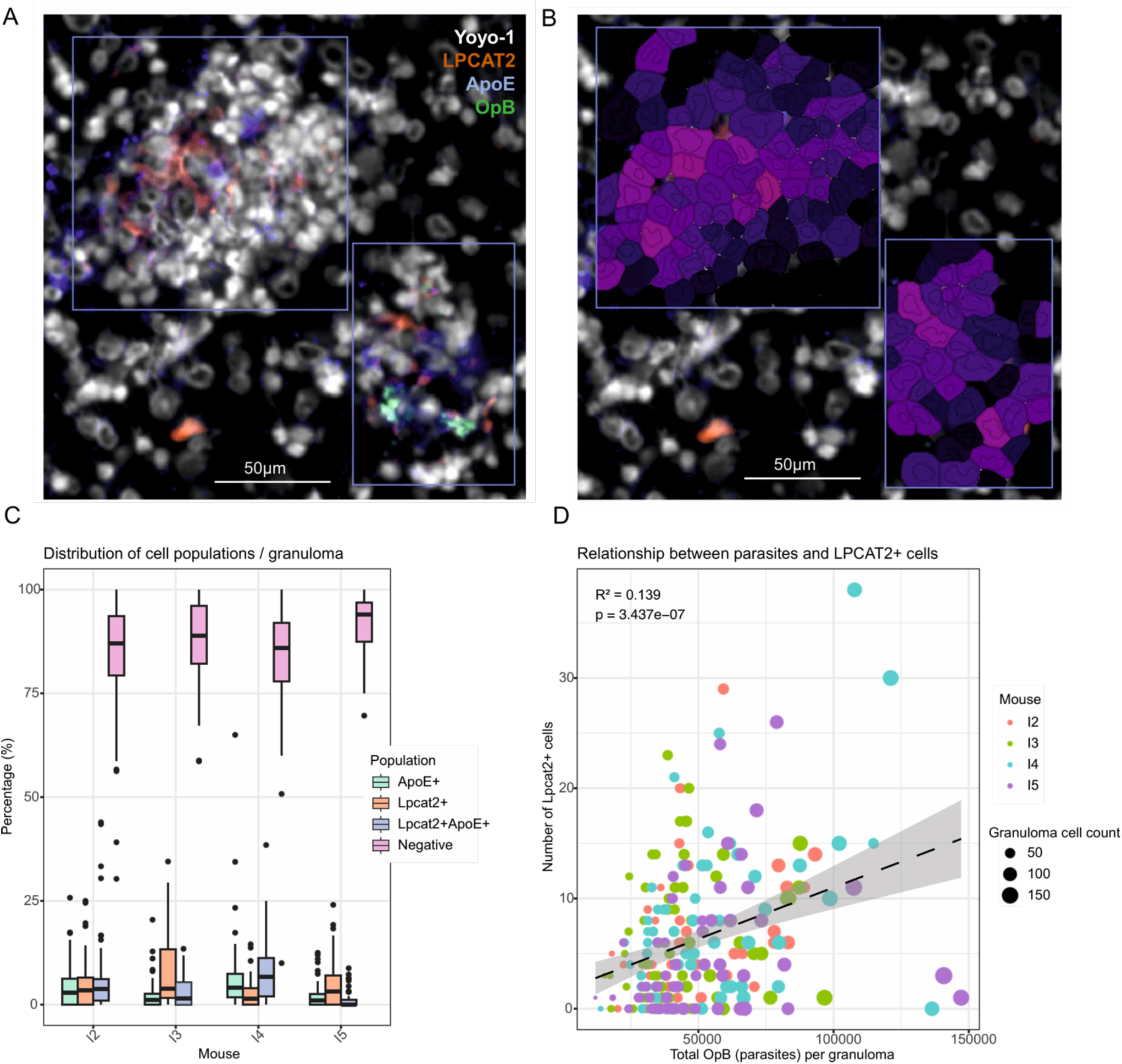
LPCAT2 protein in granulomas. **A)** Immunohistochemistry image showing nucleus counter stained with Yoyo-1 along with antibody staining with LPCAT2 (CF750), ApoE (AF594) and OpB(AF647) in naïve and **B**) infected mouse (representative). **C**) Exemplar cell segmentation based on Yoyo-1 staining and coloured on mean LPCAT2 intensity per pixel within cells (dark purple to light purple represents low to high intensity) **D**) Bar plots showing percentage of single positive (ApoE^+^ or Lpcat2^+^), double positive (Lpcat2^+^ApoE^+^) or negative cells per granuloma. Total cells; n=15,701. Box represent the interquartile range (IQR) with line representing median and whiskers show 1.5*IQR on either side **E**) Scatter plot between LPCAT2^+^ (sum of Lpcat2^+^) and parasitea (sum of OpB staining) per granuloma. Total granulomas; g=243.

### LPCAT2 is associated with macrophage activation in vivo

We used ex vivo cell sorting followed by proteomics, to further analyze the nature of LPCAT2+ macrophages within the granuloma. As LPCAT2 is expressed intracellularly, we used SIRPA / CD172 as a surrogate cell surface marker, given that the *Sirpa* transcript was correlated with granuloma spots (Figure **4G**) and macrophages with a high LPCAT2/PL remodelling score (Figure **3J**). We sorted gated CD3^-^CD11b^+^F4/80^+^ cells as CD172^hi^ (68%) and CD172^lo^ (24%) (Figure **7A** and **B**) and applied data independent acquisition proteomics to each population. As expected, sorted CD172^hi^ cells had significantly higher abundance of CD172 and LPCAT2 compared to CD172^lo^ cells. ApoE expression was not significantly different, suggesting each population of macrophages contained similar numbers of Kupffer cells (Figure **7C**). Next, we compared CD172^hi^ and CD172^lo^ cells by differential protein expression analysis. The top 5 proteins (by fold change) found in CD172^hi^ cells were TRAPPC14, WWP2, TMEM41B, CCDC127 and TMEM260 (Figure **7D**). The CD172lo cells were enriched for MCPT8, PVALB, CALM3, SDR39U1 and KRT23 (Figure **7D**).

**Figure 7:**
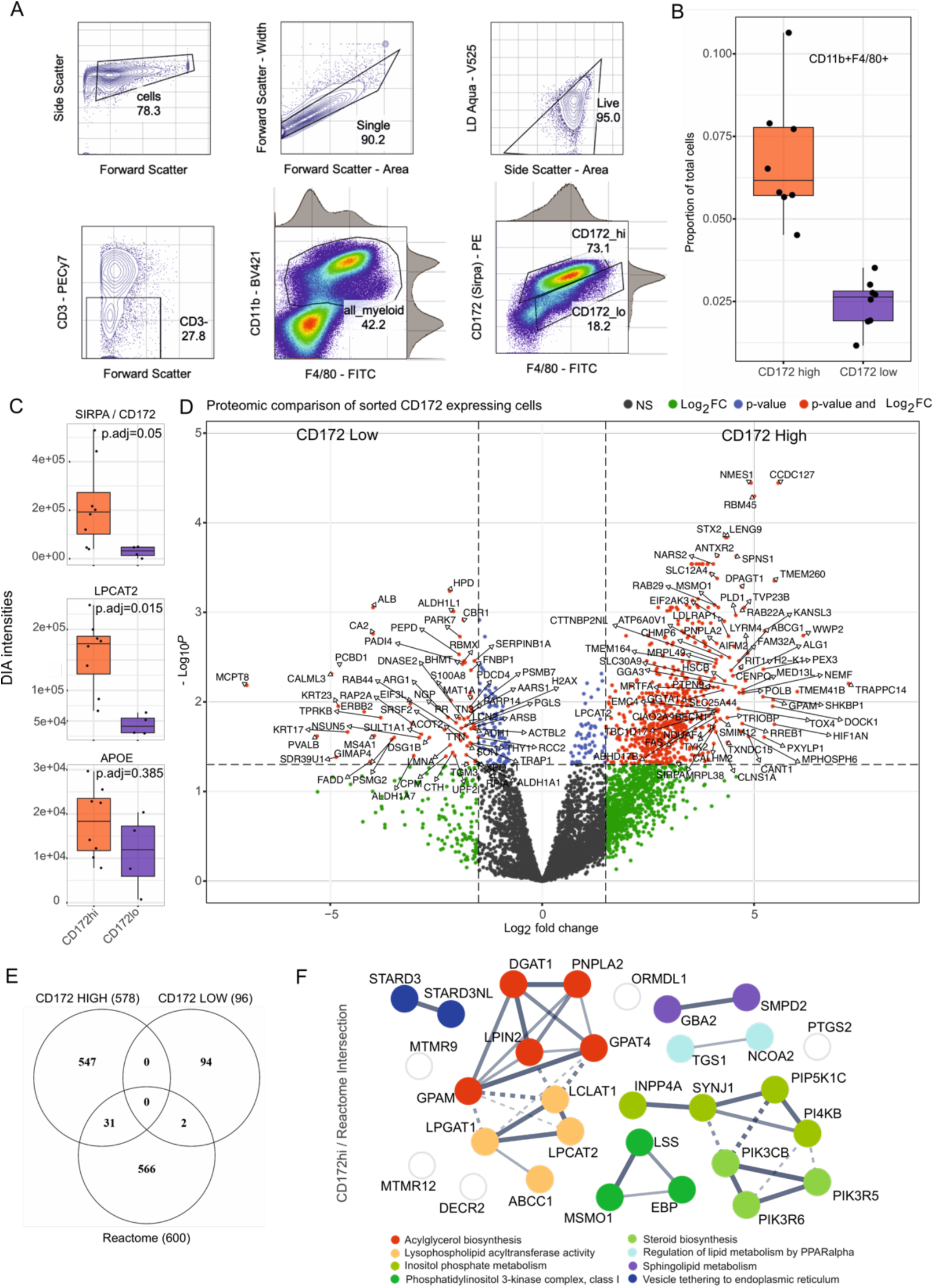
Proteomic analysis of LPCAT2 high/lo macrophages. **A)** Gating strategy to show cells selected using forward and side scatter, single cells using forward and side scatter, live cells, CD3- and then CD11b+F4/80+ cells selected as those expressing high CD172 or not **B**) Box and whiskers plot showing the spread of proportion of CD172hi versus CD172lo cells as described in A **C**) Proteomic peptide intensity between samples (n=8 for CD172^hi^; n=5 for CD172^lo^) for SIRP-α/CD172, LPCAT2 and APOE. **D**) Volcano plot showing proteomic differences between CD172 high versus CD172 low macrophages. **E**) Venn diagram showing the intersection of Reactome Lipid metabolism genes for Mus musculus with top proteins expressed by CD172 high and CD172 low macrophages. **F**) Proteins identified in E Genes identified in D are presented as a network of interactions as on STRINGDB and coloured by MCL clustering and functions labelled. Edge thickness represents interaction confidence and dotted lines indicate cluster boundaries.

GSEA analysis of all proteins upregulated (log_2_FC>2 ; FDR 5%) in the CD172^hi^ cells identified mitochondrial translation and Tnfr1-mediated signalling (known to maintain inflammatory programming^28^) as enriched pathways (**Supplementary Table CD172hi_enrichment**). CD172hi cells also expressed higher levels of CD86, whereas CD172lo cells expressed ARG1, suggesting differing states of activation between macrophages expressing physiologically higher or lower levels of LPCAT2 (Figure **7D**). Finally, we looked at proteins intersecting the Reactome Metabolism of Lipids. ∼5% of proteins from CD172^hi^ cells intersected this pathway (Figure **7E**) with pathway analysis indicating increased acylglycerol biosynthesis, and lysophoshpholipid acyltransferase activity, inositol phosphate metabolism, phosphatidylinositol 3-kinase complex activity, steroid biosynthesis, PPAR regulation, sphingolipid metabolism and vesicle trafficking (Figure **7F**).

Collectively, our multimodal data analysis indicates selective lipid mediated cell membrane re-modelling in ApoeHi_Kupffer cells and Lyz2Hi_MoMac at the heart of the granulomatous response to *L. donovani* infection.

## Discussion

In this study, we demonstrate that LPCAT2-mediated phospholipid remodeling is a defining feature of hepatic granulomas during *Leishmania donovani* infection. Through an integrated analysis combining spatial transcriptomics, lipidomics, and proteomics, we report three major findings: First, LPCAT2 expression is upregulated in macrophages within granulomas and weakly correlates with parasite burden. Second, granulomas show distinct accumulation of lysophosphatidylcholines (LPCs), the key substrates for LPCAT2, suggesting active phospholipid remodeling in these inflammatory microenvironments. Third, macrophages with high LPCAT2 expression display a pro-inflammatory phenotype characterized by their transcriptomic and proteomic signatures.

In our analysis, we define three main populations of monocytes and macrophages in the liver of infected mice. ApoeHi_Kupffer cells have abundant mRNA for *Adgre1^+^* (F4/80), *Marco*, *Clec4f* and multiple complement components and most likely represent liver resident embryonically derived Kupffer cells^29^. Of note, ApoE expression is associated with efflux of cellular cholesterol in cholesterol-loaded macrophages^30,31^ and cholesterol depletion may favour parasite survival through indirect effects on antigen presentation^32^ or direct effects on parasite survival^33^ Lyz2Hi_MoMac have abundant *Ccr2*^+^ mRNA but little *Ly62c,* and given their increased representation in infected compared to naïve mice, these likely represent monocyte-derived macrophages, that may eventually transition to a Kupffer cell phenotype^29,34,35^ These Lyz2Hi_MoMac however, have a mixed composition in terms of activation markers, expressing *Chil3* (or Ym1, Figure **2E**) known to promote Th2 immunity^36^ as well as *Tnf*, *Tlr2* and *Cd86* (Figure **S3E**). This mixed phenotype is perhaps not surprising given the highly heterogeneous cytokine profile associated with hepatic *L. donovani* infection^37^. Whether this further reflects subtleties in microenvironment or activation-induced heterogeneity in macrophages^38^ remains to be determined. Finally, we identified a smaller population of monocytes with abundant *Ly6c2* but low levels of *Ccr2* mRNA. Of these three populations, only ApoeHi_Kupffer cells and Lyz2Hi_MoMac expressed LPCAT2.

Analysis of lipid metabolism in these two macrophage populations indicated that they were enriched for SM metabolism, AA metabolism and acyl-chain remodelling of PCs via *Lpcat2* (Figure **3B-C**). LPCAT2 has been shown translocate to membrane lipid rafts and promote inflammatory gene expression^39^. LPCAT2 also participates in the Lands Cycle that re-acylates LysoPCs capable of incorporating polyunsaturated fatty acids^22^ such as arachidonic acid (AA) into the membrane that can lead to downstream processing of lipid mediators of inflammation^25^. By creating a LPCAT2/PL remodelling score (*Lpcat2, Pla2g4a, Pla2g15, Pla2g7*) we found both populations contained high scoring cells in comparable proportions. LPCAT2/PL remodelling score also positively correlated with a pro-inflammatory score.

Integrating data across modalities, we showed that the substrates for LPCAT2, lysoPCs as identified by mass spectrometry imaging are associated to immune granulomas. Palmitoyl-lysoPC(16:0), lysoPC(17:0), stearoyl-lysoPC(18:0), linoleoyl-lysoPC(18:2) and arachidonoyl-lysoPC(20:4) were all detected. Palmitoyl and arachidonyl-lysoPCs are known to promote Cox-2 expression^40^, a key enzyme for AA-led inflammation, and *Ptgs2* (Cox-2) and *Tbxas1*, involved in downstream AA metabolism, were upregulated in macrophages and localized to granulomas. These data are in accord with our previous observation of enrichment of arachidonic acid (AA)-containing lipids in *Leishmania* granulomas^20^.

We confirmed the expression of LPCAT2 by *ex vivo* proteomic analysis of macrophages, using SIRPA / CD172 as a surrogate for LPCAT2 expression. In line with *in vitro* findings of LPCAT2 having a pro-inflammatory effect on macrophages^39^ we found that CD172^hi^ cells were enriched for pro-inflammatory pathways However, macrophage phenotype with respect to lipid composition is clearly complex and likely involves elaborate lipid remodelling by lysophospholipid acyltransferases not limited to LPCAT2. For example, multiple ether-linked PCs that have been linked to be sensitive for ferroptosis^41^ such as PC(O-36:3), PC(O-34:2) were also found in granulomas and CD172^hi^ macrophages also expressed PEX3) a peroxisomal biogenesis factor, essential for ether-linked PC synthesis^42^.

Finally, our data have implications for the tissue- and context-specific regulation of lipid metabolisms by LPCAT enzymes. In lung homeostasis, acyl chain remodeling of phosopholipids essential for pulmonary function occurs through LPCAT1^43^, and this pathway is enriched in lung disease^44^. In liver homeostasis, LPCAT3 is important for lipid homeostasis^27^, and in our study was upregulated in infected mice in spots rich in hepatocytes. However, within immune granulomas LPCAT2 is highly expressed in both resident and recruited macrophages. LPCATs and their substrates LPCs have high therapeutic potential, being implicated in conditions from pain^45^ to cancer metastasis^46^. They also function as lyso-platelet activating factor (PAF) acyltransferases, synthesizing PAF in a range of proinflammatory diseases, and selective LPCAT2 inhibitors have recently been identified^47^ opening the door for future therapeutic manipulation. However, in the context of leishmaniasis, where LPCATs maintain a balance between the pro-(PAF-mediated) and anti-(prostaglandin-mediated) inflammatory activities, the outcome of LPCAT2 inhibition remains to be determined.

Our study has some limitations. One shortcoming is that whilst our spatial resolution is 55μm, spots have a centre-to-centre distance of 100μm resulting in voids where measurement does not occur. Similarly, some spots on the periphery of granulomas may include parenchymal cells. These issues are partly mitigated, however, by the sampling of large numbers of granulomas and their lack of a defined shape / compartmentalization. The marriage between single cell and spatial information is also largely probabilistic and not exact, but we have used orthogonal approaches to cross validate key findings. Finally, although we observed a weak but significant correlation between LPCAT2 expression and granuloma parasite load, it was not possible to definitively link intracellular parasitism with the changes observed. Although parasites were observed in some granulomas, thin tissue sections do not sample the entire granuloma volume and may underestimate parasite load.

Given that parasites are largely found in embryonically derived Kupffer cells^35^ but we found that both ApoeHi_Kupffer cells and Lyz2Hi_MoMac regulate LPCAT2, it is likely that many of the signals governing macrophage activation state and LPCAT2 expression are cell-extrinsic and derived from other leucocytes in the granuloma microenvironment.

In conclusion, our study provides a comprehensive map of the immunometabolic landscape within *L. donovani-*induced granulomas. The identification of LPCAT2 as a key player in the macrophage response during granulomatous inflammation brings forth new avenues for understanding and potentially modulating the immune response in leishmaniasis and possibly other granulomatous diseases. Future studies should focus on elucidating the functional consequences of lipid remodelling in granulomas and exploring the therapeutic potential of targeting these pathways.

## Materials & Methods

### Ethics statement

All animal studies were performed under UK Home Office Licence (PP0326977) and were approved by the Animal Welfare and Ethical Review Board of the University of York.

### Animals and Infection Model

Adult female C57BL/6J mice were maintained and bred at the University of York, UK. The mice were housed in ventilated cages under specified pathogen-free conditions with access to food and water *ad libitum*. Identification of the mouse colony was confirmed through genetic profiling of microsatellite markers, verifying C57BL/6J genetic background with minor variations at 3 markers. Healthy (6-8 week old females) were either infected on day 0 intravenously with 3×10^7^ amastigotes of the Ethiopian LV9 strain of *Leishmania donovani* to establish visceral infection or left naïve. At 28 days post-infection, mice were euthanized by CO_2_ inhalation followed by cervical dislocation to model peak granulomatous inflammation in the liver. Livers were harvested after euthanization and the major lobe cut and snap-frozen in liquid nitrogen while bathed in isopenane and then kept in -80 degrees until used. The remaining liver was harvested in 2% fetal calf serum (FCS) in 1x PBS and immediately processed into a single cell-suspension. Age-matched naïve female mice from the same colony were used as baseline controls for liver tissue. Individual mice were used as experimental units unless stated in the figure legends. Aged-matched mice were randomly (coin-toss) allocated to groups, and no blinding or exclusion was employed due to clear experimental conditions (infected vs. naive mice). Specific *a priori* effect size calculation for the multi-modal analysis was not performed. A total of 25 mice were used across 3 independent experiments: Spatial transcriptomics and mass-spectrometry based lipidomics (n=8; 4 infected and 4 naïve), single-cell RNA-seq (n=12; 6 infected (3 matched mice from spatial transcriptomics and 3 from an independent experiment) mice plus 2 pools of 3 naïve mice each), and flow-sorted proteomics (n=8 infected mice, from which CD172^hi^ macrophages [n=8 samples] and CD172^lo^ macrophages [n=5 samples] were obtained). Due to low cell recovery post-sort in the CD172^lo^ fraction, these could not be processed for proteomics leading to 3 unmatched CD172^hi^ animals.

### Liver dissociation and single-cell suspension

Liver tissue in 2%FCS in 1xPBS was sliced coarsely using a scalpel and then kept at 37 degrees in an incubator with 5% Co2 in HBSS with collagenase and Dnase for 20 minutes. Ice cold FCS containing RPMI was used to stop the enzymatic digestion by collagenase. Tissue was then passed through a 100μm cell strainer by mashing the tissue with the plunger of a 5ml syringe. Upon completion, cells were suspended in ACK lysing buffer for 5 minutes. Cells were washed and then were centrifuged for 15 minutes without breaks in 33% Percoll to separate the hepatocytes.

Hepatocytes formed a brown ring on top which was then discarded by decanting. The remaining cell population was washed twice and then re-suspended in PBS containing 0.05% BSA as per 10x instructions. Cells were either used for single-cell RNA seq or flow cytometry.

### Cryo-sectioning and histopathology

Frozen sections were cut in a Leica cryostat while maintaining them on a pedestal of OCT without allowing OCT to touch the tissue itself. 7μm sections were cut onto PEN membrane slides, 10x Visium spatial transcriptomics gene expression slides and onto superfrost glass slides. The order of the sections was always, super frost 1, PEN membrane 1, Visium 1, Pen Membrane 2, superfrost 2 for each mouse. PEN membrane 1 and 2 were used for mass spectrometry-based imaging in positive and negative ionization mode measurements. The Visium slide was used for spatial transcriptomics.

### Mass spectrometry imaging of lipids and lipid identification experiments

All PEN microdissection membrane slides were dried for 15 min under vacuum and fiducial markers (Tipp-Ex, BIC, France) were applied to the slides before matrix application. The norharmane matrix solution at concentration of 7 mg/ml was prepared in 2:1 chloroform: methanol (v:v). An automated TM sprayer (HTX Technologies, Chapel Hill, NC, U.S.A.) was used for matrix application. In short, 12 layers were sprayed homogeneously at a fixed flow rate of 0.12 ml/min with the spray nozzle temperature at 30°C combined with drying time between every layer for 30 seconds. After matrix application, lipidomics mass spectrometry imaging experiments were performed using a timsTOF fleX (Bruker Daltonik, Bremen, Germany) at the mass range of *m/z* 300–1800 with laser frequency of 5000 Hz and 50 laser shots accumulated per pixel in both negative and positive ionization modes. A total of 8 liver sections from 8 mice were measured with a pixel size of 20*20 µm^2^ for each polarity.

For the lipid identification, data-dependent analysis (DDA) imaging experiments were performed on 2 representative mouse liver sections (one naïve mouse and one infected mouse) using an LTQ Orbitrap Elite mass spectrometer (Thermo Fisher Scientific, Bremen, Germany) as previously described by Ellis *et al*. In short, a 240,000-mass resolution full-scan FTMS (Orbitrap) analysis was conducted in the mass range between *m/z* 300–1800 combined with stage step size of 25 µm (horizontal) and 50 µm (vertical). Subsequently, DDA IT-MS/MS (ion trap) scans were performed at adjacent 25-µm positions.

Automated lipid identification was performed using LipoStarMSI^48^ (edition 2023) incorporated with the publically available Lipid Maps database. Precursor *m/z* tolerance was 0.00 Da ± 3.00 ppm and MS/MS *m/z* tolerance was 0.25 Da ± 0.00 ppm. The considered ion adduct types are [M+H]^+^, [M+Na]^+^, and [M+K]^+^ for the positive ion mode and [M-H]^-^ for the negative ion mode.

### Histological characterization & annotations for MSI

After MALDI-MSI measurement, the matrix was removed with 100% ethanol prior to hematoxylin staining. All slides were firstly rinsed in a graded ethanol series (100%, 96%, 70%, 2 min each), followed by immersing in distilled water for 2 min. Subsequently, the sections were stained with 0.1% Gill’s Hematoxylin (Merck, Darmstadt, Germany) for 3 min, followed by rinsing in running tap water for 3 min and dehydrated in a graded ethanol series (70%, 2*96%, 2*100%, 2 min each).

Finally, all slides were washed in 100% xylene for 2 min and not covered with cover slip. After drying overnight in desiccator, the hematoxylin stained slides were scanned using Leica Aperio CS2 slide scanner at 20x magnification (Leica Microsystems, Nussloch, Germany). All scanned high-resolution hematoxylin stained images were coregistered in FlexImaging v5.0 (Bruker Daltonik GmbH, Bremen, Germany) to the MSI data using the previously applied fiducial markers.

The acquired MSI data and coregistered hematoxylin stained images were imported into the mass spectrometry imaging software SCiLS Lab (version 2023b) for both polarity experiments separately without baseline removal and automatic resampling. Pixels of the negative and positive polarity datasets were normalized to their root-mean-square or total-ion-count values, respectively. Peak picking was done on the average spectrum of every dataset using mMass and the following parameters: S/N>5, relative intensity threshold > 0.2%, picking height 90%, and de-isotiping with mass and intensity tolerances of *m/z* 0.1 and 70%, respectively. Peak lists were re-imported with peak area as interval processing mode and a fixed interval width of 6 mDa and 10 mDa for the positive and negative polarity data, respectively, into SCiLS Lab.

### Spatial Transcriptomics

Sections from uninfected and *L. donovani* infected mice (n=4 per group, 8 total) attached to 10x Genomics Visium slides were processed using the Visium Spatial Gene Expression workflow as per manufacturer’s instructions for fresh frozen tissue. Sections were fixed in pre-chilled methanol and then stained in hematoxylin for 5 minutes, bluing buffer for 1 minute and finally stained in eosin Y for 30 seconds. Coverslips were added using glycerol containing RNase inhibitor and the slides imaged using a Zeiss axioscan slidescanner. Coverslip was removed in MQ water and then the slides were placed in a gene expression cassette. H&E stained tissue sections were then permeabilized by proteinase K then reverse transcribed to synthesize cDNA directly on the slide. Second strand cDNA synthesis was performed for 25 minutes followed by cDNA denaturation for 15 minutes. The optimal number of amplification cycles was determined by qPCR then cDNA was amplified. Amplified cDNA was cleaned up with SPRIselect beads and quantified.

Fragmentation, end repair, A-tailing and adaptor ligation were performed to construct sequencing libraries. Post-ligation cleanup, sample index PCR and double-sided SPRIsize selection were conducted. Final libraries were quality controlled on an Agilent Bioanalyzer. cDNA libraries were constructed per kit instructions. Quantified libraries were sequenced on an Illumina NovaSeq 6000 to generate spatially barcoded, transcriptomic sequencing data aligned to H&E reference images Raw sequencing data was aligned to the mouse mm10 genome using 10x spaceranger (1.3.0) software.

H&E images were aligned to fiducial markers on each slide using the Loupe browser. Tissue regions were manually selected in Loupe and coordinate JSON files created. These JSON files were input into spaceranger count() alongside sequencing data to generate spatially-resolved gene counts. Raw counts were loaded as a .h5 file in Seurat ^49^ for analysis.

### Image registration

To assist image registration between MSI and spatial transcriptomics (Visium) data, segmentation images were first created for the MSI data of infected mice in SCiLS lab using bi-secting k-means with correlation distance. Subsequently, the peak picked and normalized MSI data, its co-registered H&E image (H&E_MSI_), and existing segmentation images were read-out from SCiLS Lab directly to R (version 4.1.0) using the API package “SCiLSLabClient”. From there the data was transferred to MATLAB (version 2018a) via a .mat file using the R package “R.matlab”. In MATLAB naïve mouse MSI datasets were segmented using k-means with *k*=3. Finally, the Visium data and its associated H&E images (H&E_Visium_) were also imported into MATLAB where the registration between MSI and Visium has been performed in a 2-step approach: First, a manual, coarse control point-based affine registration has been performed between H&E_MSI_ and H&E_Visium_. Since the MSI data is also registered with a certain error to its H&E_MSI_, the MSI data was in a second step directly registered to the H&E_Visium_ using the MSI segmentation image which shows morphological structures that can be found back in H&E_Visium_. This step can be considered a refinement of the first registration and was also done manually using control points with affine or with piece-wise linear transformation, depending on the individual sample pairs’ needs. This process was repeated for all individual datasets and resulted in the assignment of approximately 4–6 MSI pixels to the 55 μm-sized Visium spots. Pixels failing co-registration between MSI and Visium were excluded from downstream analysis but this did not lead to animals being excluded from the study.

### Immunohistochemistry

Frozen liver sections (7μm) from naive and infected mice were cut using a Leica cryostat, placed onto superfrost slides. Sections were fixed in ice-cold acetone for 10 minutes followed by two washes (5 minutes each, with agitation) in wash buffer (PBS with 0.05% BSA, w/v). Hydrophobic barriers were then applied around each tissue section. Sections were blocked for 30 minutes at room temperature (RT) using dilution buffer comprising 5% donkey serum diluted in wash buffer supplemented with 0.1% Triton X-100. Primary antibodies targeting OpB (in-house; sheep IgG, 10 µg/ml) and LPCAT2 (Invitrogen PA5-101406; rabbit IgG, 1:500 dilution) were applied simultaneously in blocking buffer for 1 hour at RT. Secondary antibodies donkey anti-sheep IgG AF647 (1:750 dilution) and donkey anti-rabbit IgG CF750 (1:500 dilution) were subsequently applied for 1 hour at RT, prepared in wash buffer. Next, directly conjugated anti-Apoe antibody (rabbit monoclonal clone EPR19392, AF594 conjugate, Abcam ab310350, 1:500 dilution) and nuclear stain Yoyo-1 (0.2 µM, 1:500 dilution from 100 µM stock solution) were applied together in wash buffer for 1 hour at RT. Finally, sections were mounted using Prolong Gold mounting media and incubated at 4°C in the dark for 24 hours prior to imaging.

### Single-cell RNA Sequencing

Single cells were resuspended at a concentration of approximately 1,000 cells per microliter. Single cell RNA sequencing library preparation was then carried out using the Chromium Next GEM Single Cell 3’ Kit v3.1 from 10X Genomics (Pleasanton, CA), following the manufacturer’s CG000315 Rev A user guide. For library preparation, a target of 10,000 cells was captured per sample. Sequencing libraries were prepared as outlined in the 10X Genomics user guide and sequenced on the Illumina NovaSeq 6000 system (San Diego, CA) to achieve a minimum sequencing depth of approximately 20,000 reads per cell.

### Single-cell RNA data pre-processing

The raw FASTQ files generated from sequencing were aligned to the mouse reference genome (mm10) using the Cell Ranger software (version 6.1.0) from 10x Genomics. Cell Ranger Count was used to align reads and generate gene-barcode matrices for each library. The resulting .h5 files were imported into the R package Seurat (version 4.3.0) for quality control and downstream analysis. Low quality cells were filtered out by removing any cells with greater than 10% mitochondrial reads. The data was visualized using Seurat, plotting cells based on feature-feature relationships to identify cell populations and subset the data accordingly. Potential library construction artifacts were removed by dropping read counts for transcripts Gm42418 and AY036118. Data was normalized using SCTransform, implementing a gamma-Poisson generalized linear model (glmGamPoi)^50^ to regress out variation due to mitochondrial reads, ribosomal genes, total RNA content, and unique feature counts. This normalized, filtered gene count matrix was used for subsequent analyses. Similar approaches for quality control of the spatial data were performed.

### Spatial Data Analysis

The positive and negative matrices per Visium pixel as generated from image co-registration resulted in an averaged matrix of MSI intensities [Vm x Mi] with species *m/z* measure in the positive and a second matrix [Vm x Mj] in the negative ionization mode Vm refers to ‘m’ spatial barcodes measured by the 10x Visium platform. The .h5 file from spatial transcriptomics also contained a matrix [Vm x Tk] which represented 1..k transcripts measured across 1..m barcoded spots from now referred to as the transcript matrix. At first, the matrices from positive and negative ionization measurements were combined i.e. a full join was used to obtain all Vn spots measured resulting in a matrix [[Vm x Mi] | [Vm x Mj]] or referred to as the MSI matrix [Vm x Mi+j].

Initially, each of the matrices were analysed separately for variable features and a principal components (PC) analysis was performed separately. The reduced PC space was then used as an input for finding clusters using Louvain clustering and as an input for t-SNE. Each Vm was then coloured by its cluster identity and visualized in the 2-dimensional t-SNE space. When the PCs were deduced from the transcript matrix the colour coded clustered were referred to as RNA clusters, while when the PCs were deduced from the lipid abundance intensities from the MSI matrix the cluster labelling was labelled as Lipid clusters.

For sub-clustering only spots form three clusters (i.e. RNA_0/4/7) was subsetted. The sub-clustering was done using transcript data only and MSI data was retained as metadata. Single-cell RNA seq matrices were clustered using a reduced PC space and then clusters were checked for types by looking gene expression profiles by calculating DE per cluster identity by taking one versus all approach. Genes were then looked for canonical markers and cell types assigned.

The cell type information along with the single cell expression dataset and the Visium dataset was used as input for Cell2Location (v0.1) to model expression in single cells and then use that to deconvolute cell type abundances in space (Visium). The 5^th^ quantile of the predicted abundances was used for all downstream analysis. Since the abundances were associated for each Vn this data is then easily integrated across the dataset. Correlations between lipid intensities and predicted cellular abundances were calculated using Pearson’s correlation and the R function corr. Only statistically significant correlations were reported. Cell type abundances were compared between sub-clusters using a Kruskal-Wallis test with multiple comparisons to check for difference between groups. P-value of significance was set at 0.05.

### FACS Sorted Proteomics

Single-cell suspensions were stained in PBS with 0.05% BSA, using fluorescent antibodies against CD3, CD11b, F4/80, and CD172 (SIRPA). Macrophage populations (CD3–CD11b+F4/80+) were sorted on a Beckman Coulter MoFlo Astrios, separating cells based on CD172 expression into CD172hi and CD172lo subsets. Sorted cell populations were diluted 1:1 with aqueous 10% (v:v) sodium dodecyl sulfate, 100 mM triethylammonium bicarbonate (TEAB). Protein was reduced with 5.7 mM tris (2-carboxyethyl) phosphine and heating to 55°C for 15 minutes before alkylation with 22.7 mM methyl methanethiosulfonate at room temperature for 10 minutes. Protein was acidified with 6.5 µl of aqueous 27.5 % (v:v) phosphoric acid then precipitated with dilution seven-fold into 100 mM TEAB90% (v:v) methanol. Precipitated protein was captured on S-trap (Profiti – C0-micro) and washed five times with 165 µl 100 mM TEAB 90% (v:v) methanol before digesting with the addition of 20 µl 0.1µg/µl Promega Trypsin/Lys-C mix (V5071) in aqueous 50 mM TEAB and incubation at 47°C on hotplate for 2h. Peptides were recovered from S-trap by spinning at 4000 g for 60 s. S-traps were washed with 40 µl aqueous 0.2% (v:v) formic acid and 40 µl 50% (v:v) acetonitrile:water and washes combined with the first peptide elution. Peptide solutions were dried in a vacuum concentrator then resuspended in 20 µl aqueous 0.1% (v:v) formic acid. Peptides were loaded onto EvoTip Pure tips for nanoUPLC using an EvoSep One system. A pre-set 30SPD gradient was used with a 15 cm EvoSep C18 Performance column (15 cm x 150 μm x 1.5 μm). The nanoUPLC system was interfaced to a timsTOF HT mass spectrometer (Bruker) with a CaptiveSpray ionisation source. Positive PASEF-DIA, nanoESI-MS and MS2 spectra were acquired using Compass HyStar software (version 6.2, Bruker). Instrument source settings were: capillary voltage, 1,500 V; dry gas, 3 l/minute; dry temperature; 180°C. Spectra were acquired between m/z 100-1,700. DIA windows were set to 25 Th width between m/z 400-1201 and a TIMS range of 1/K0 0.6-1.60 V.s/cm^2^. Collision energy was interpolated between 20 eV at 0.65 V.s/cm2 to 59 eV at 1.6V.s/cm^2^.

LC-MS data, in Bruker .d format, was processed using DIA-NN (1.8.2.27) software and searched against and in-silico predicted spectral library, derived from the mouse subset of UniProt appended with common proteomic contaminants. Search criteria were set to maintain a false discovery rate (FDR) of 1%. High-precision quant-UMS^51^ was used for extraction of quantitative values within DIA-NN. Peptide-centric output in. tsv format, was pivoted to protein-centric summaries using KNIME 5.1.2 and data filtered to require protein q-values < 0.01 and a minimum of two peptides per accepted protein. Calculation of log2 fold difference and differential abundance testing was performed using limma via FragPipe-Analyst^52^. Sample minimum imputation was applied and the Hochberg and Benjamini approach was used for multiple test correction.

### Statistical assumptions

Data were analyzed with standard exploratory techniques as described under the sections *Single-cell RNA data pre-processing*, *Spatial Data Analysis*, *FACS Sorted Proteomics* and formal testing for statistical assumptions was not performed.

## Author contribution

SD, PMK and RH conceived the study. SD and JHC designed and performed the experiments. SD analysed the integrated data, created visualisations and wrote the manuscript. BB, JHC co-registered the images. SD, NSD conducted the mouse experiments, and the spatial transcriptomics. SJ and LG prepared libraries for sequencing from barcoded Visium spots. AD conducted proteomic data analysis. GC and POT provided insights and feedback. All authors edited the manuscript. PMK and RH reviewed the manuscript and supervised the work.

## Acknowledgement

The authors would like to acknowledge the help of Dr Karen Hogg and Dr Karen Hodgkinson at the University of York’s Bioscience Technology Facility I&C lab and that of Dr. Angelo Lopez and Dr. Chloë Baldreki within the University of York’s Bioscience Technology Facility MAP-lab, for assistance with proteomic sample processing and LC-MS/MS data acquisition. The York Centre of Excellence in Mass Spectrometry was created thanks to a major capital investment through Science City York, supported by Yorkshire Forward with funds from the Northern Way Initiative, and subsequent support from EPSRC (EP/K039660/1; EP/M028127/1).

## Data Availability

Spatial and single-cell transcriptomic data available on gene expression omnibus with the accession codes GSE290324 and GSE290325 respectively. All related code is available on https://github.com/jipsi/spatial_lipid_gene. Proteomic mass spectrometry data sets and results files are referenced in ProteomeXchange (PXD058137) and available to download from MassIVE (MSV000096486) [doi:10.25345/C5ST7F84G].

## Supplementary Figures

**Figure S1:**
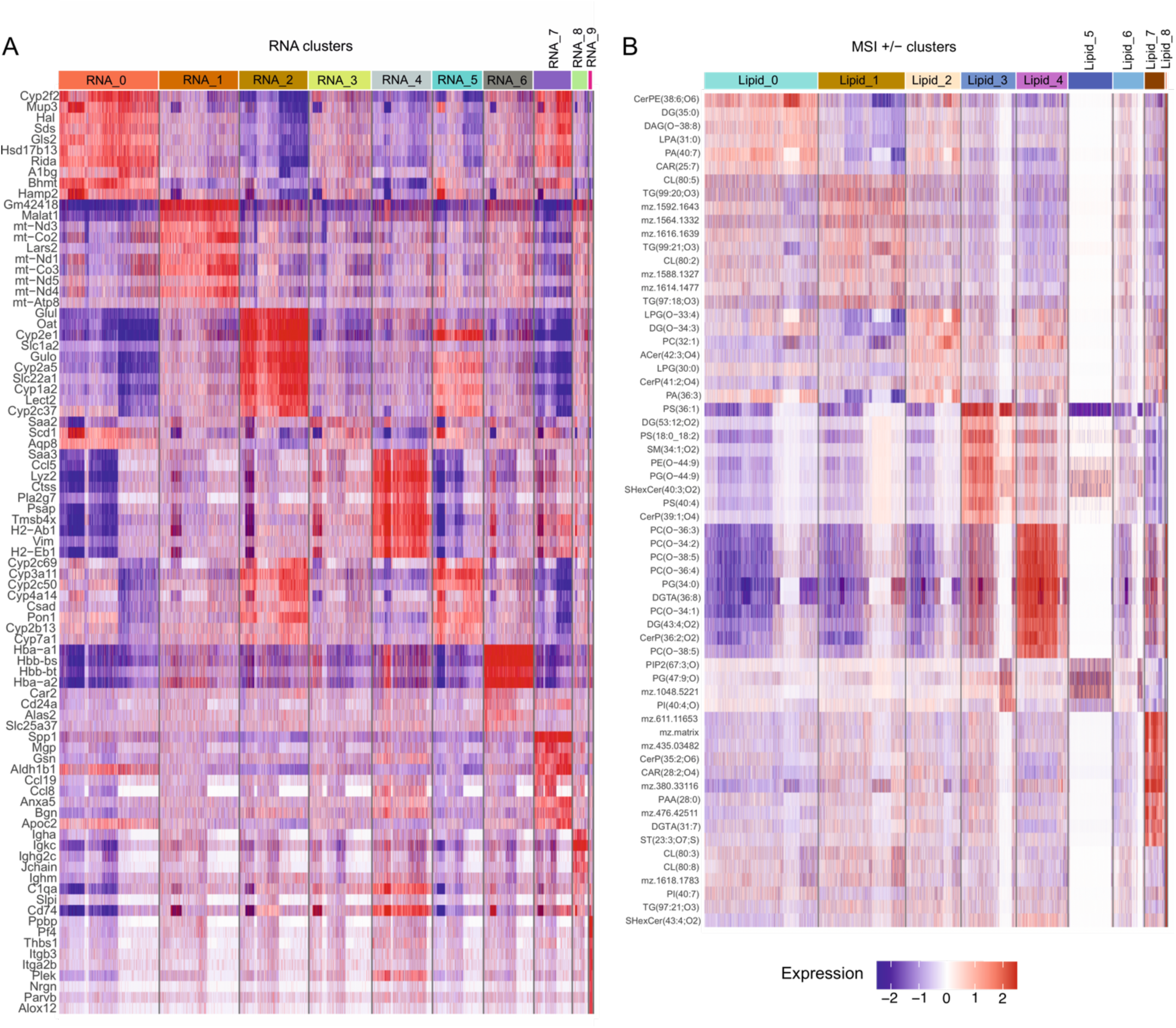
Heatmap showing differential transcription (A) and lipid species (B) between RNA_clusters and Lipid_clusters respectively

**Figure S2:**
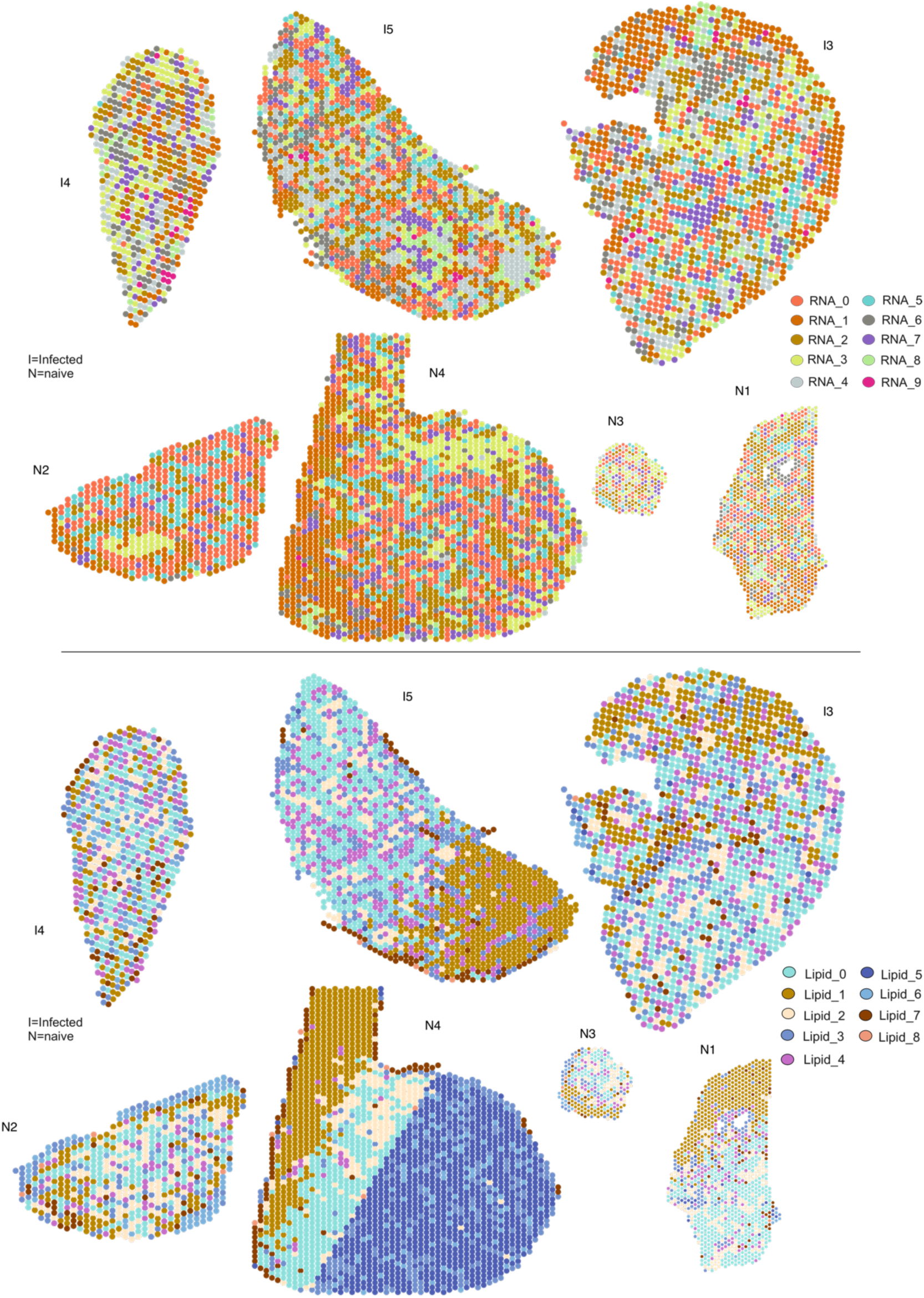
Spatial plot showing a representative infected section with spatial spots coloured by their cluster colour (RNA clusters - top and Lipid clusters - bottom)

**Figure S3:**
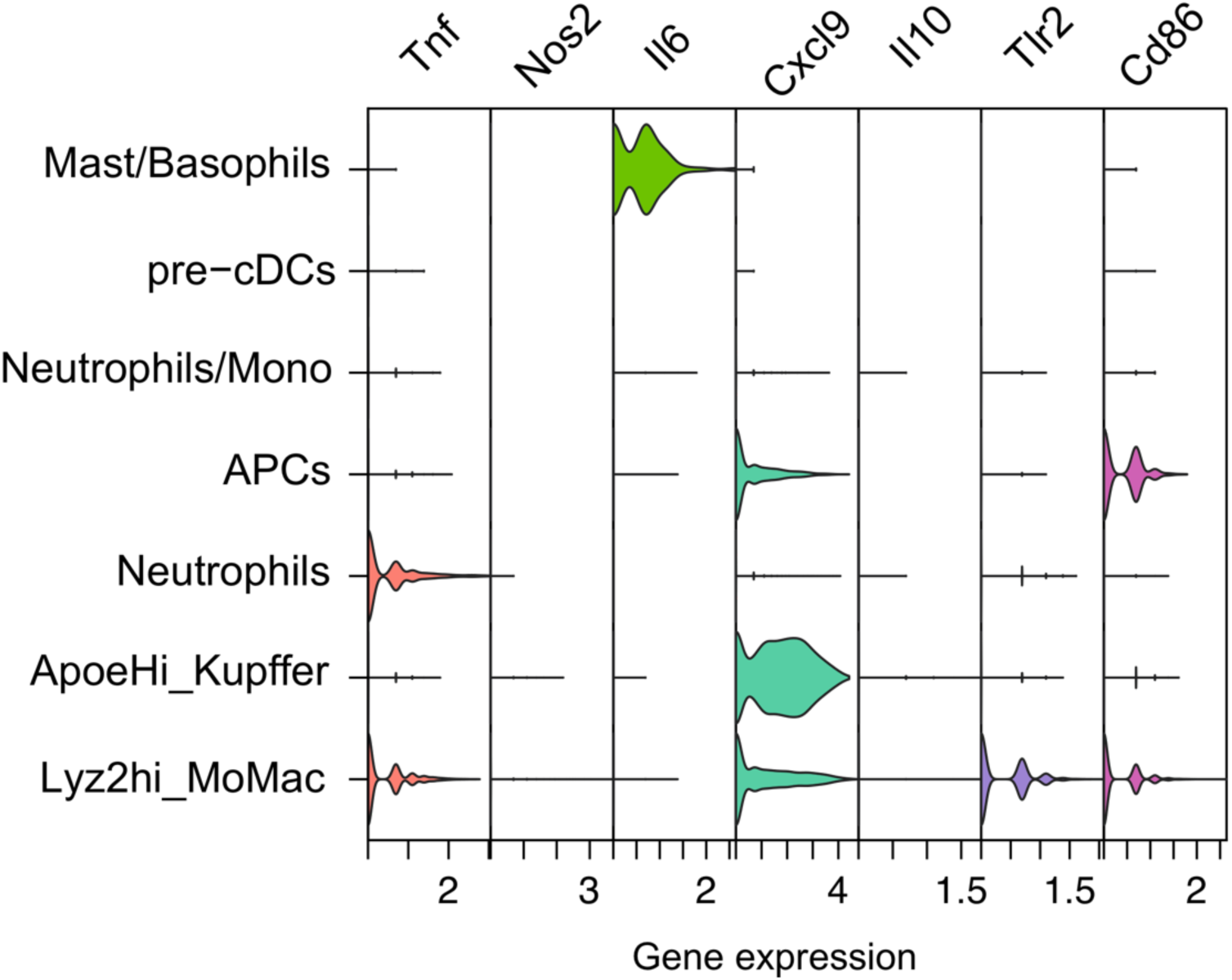
Stacked violin plot showing expression of Tnf, Nos2, Il6, Cxcl9, Il10, Tlr2, Cd86 across myeloid populations

**Figure S4:**
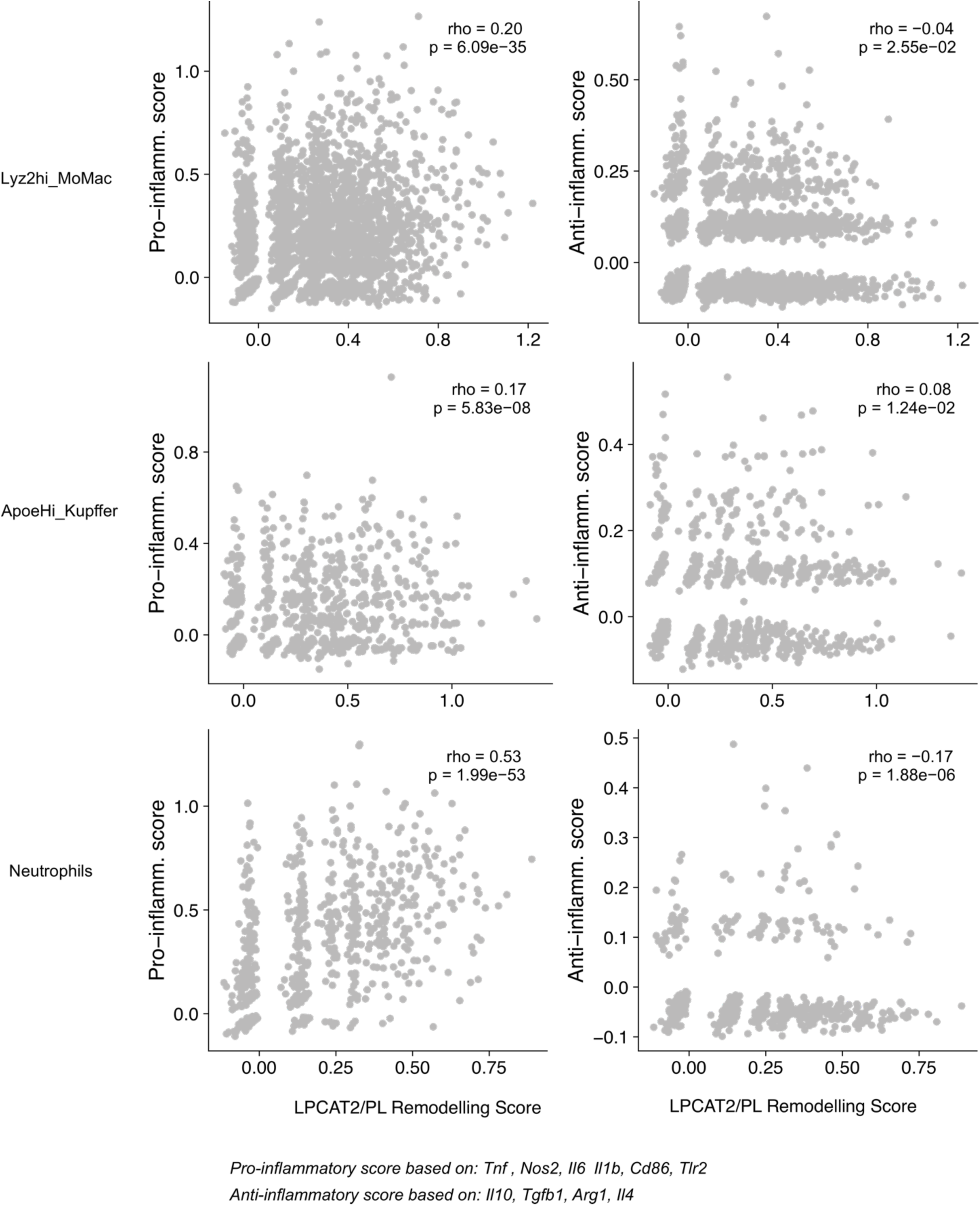
Scatter plots showing Spearman’s correlation (between LPCAT2/PL Remodelling Score and either Pro-inflammatory score (left panels) or Anti-inflammatory score (right panels), for Lyz2HiMoMac (top), apoeHi_Kupffer (middle) and Neutrophils (bottom).

**Figure S5:**
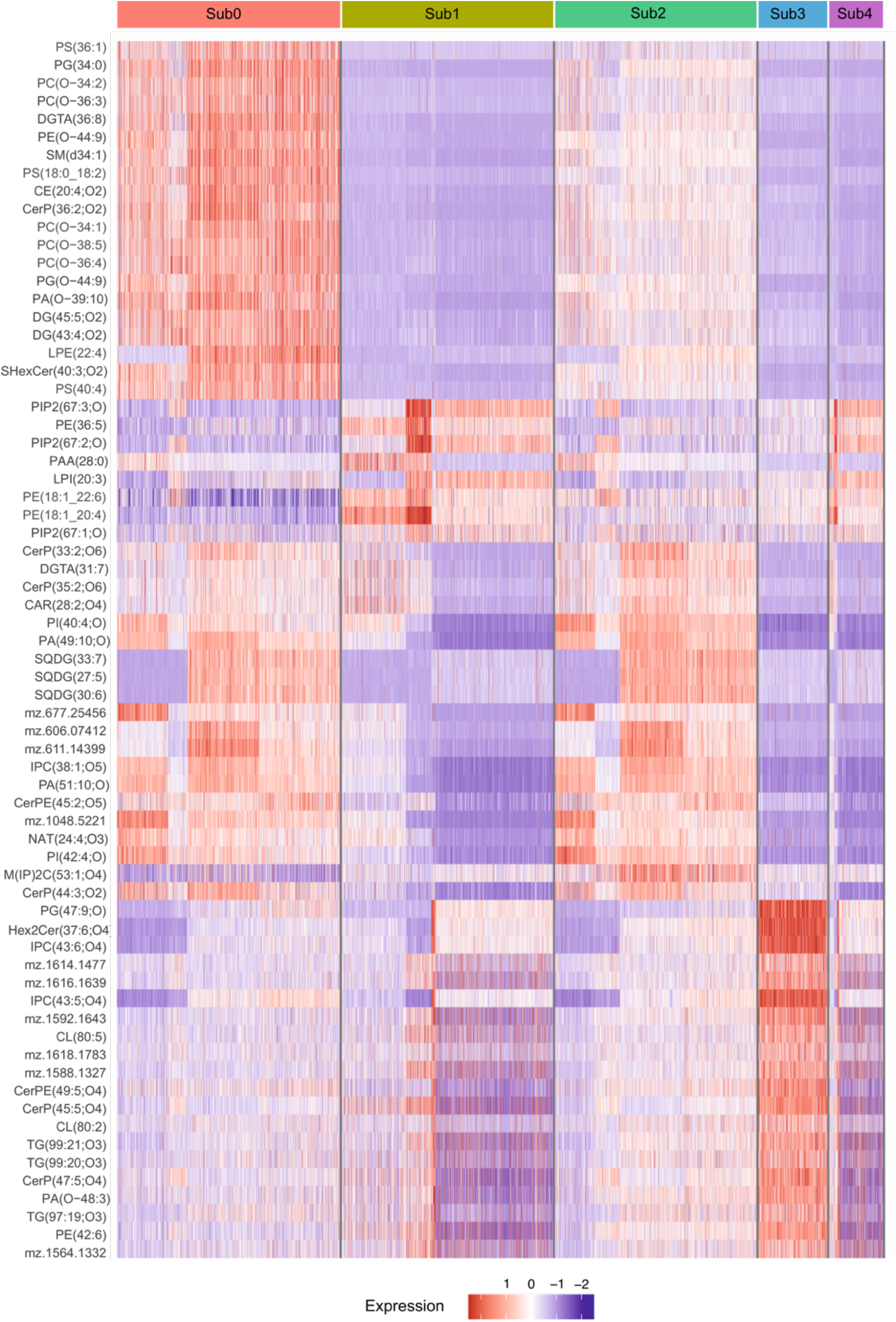
Heatmap showing differential lipid species between sub-clusters of RNA0/4/7 i.e. Sub0, Sub1, Sub2, Sub3, Sub4. Sub0 and Sub2 are groups specific to infection only and Sub0 represent immune granulomas.

